# An INS-1 β-cell proteome highlights the role of fatty acid biosynthesis in glucose-stimulated insulin secretion

**DOI:** 10.1101/2024.07.12.603204

**Authors:** Nina Stremmel, Oliver Lemke, Kathrin Textoris-Taube, Daniela Ludwig, Michael Mülleder, Julia Muenzner, Markus Ralser

## Abstract

Pancreatic beta cells secrete insulin as a response to rising glucose level, a process known as glucose-stimulated insulin secretion (GSIS). In this study, we used liquid chromatography tandem mass spectrometry and data-independent acquisition to acquire proteomes of rat pancreatic INS-1 832/13 beta cells that were short-term stimulated with glucose concentrations ranging from 0 to 20 mM, quantifying the behavior of 3703 proteins across 11 concentrations. Ensemble clustering of proteome profiles revealed unique response patterns of proteins expressed by INS-1 cells. 237 proteins, amongst them proteins associated with vesicular SNARE interactions, protein export, and pancreatic secretion showed an increase in abundance upon glucose stimulation, whilst the majority of proteins, including those associated with metabolic pathways such as glycolysis, the TCA cycle and the respiratory chain, did not respond to rising glucose concentrations. Interestingly, we observe that enzymes participating in fatty acid metabolism, responded distinctly, showing a “switch-on” response upon release of glucose starvation with no further changes in abundance upon increasing glucose levels. We speculate that increased activity of fatty acid metabolic activity might either be part of GSIS by replenishing membrane lipids required for vesicle-mediated exocytosis and/or by providing an electron sink to compensate for the increase in glucose catabolism.

**Significance of the Study:** We used high-throughput proteomics to capture comprehensive proteome changes 30 minutes post stimulation in the INS-1 832/13 beta cell line. Our study provides insights into the metabolic regulation of glucose-stimulated insulin secretion in pancreatic beta cells, specifically highlighting the early role of fatty acid biosynthesis. These findings suggest a necessary shift in focus from electrochemical to metabolic mechanisms in understanding GSIS, paving the way for future research. As the first to document proteome alterations in the initial phase of GSIS, our study furthermore documents the extent of protein abundance variability when obtaining data after short stimulation times, and therefore highlights the necessity of well-controlled study design and biological replicates. The recorded data set complements existing metabolomic and transcriptomic studies, providing a valuable resource for subsequent investigations.

## Introduction

Pancreatic islets are assemblies of endocrine cells spread throughout the pancreas. Endocrine beta cells account for the majority of islet cells in all vertebrates and secrete insulin as a response to rising blood glucose levels. Impaired insulin secretion causes severe diseases like Type-1 and Type-2 Diabetes Mellitus, which together affect around 537 million people worldwide [1]. Glucose-stimulated insulin secretion (GSIS) is highly regulated [2,3] and occurs in two phases. The triggering phase occurs during the first 20 minutes after a glucose stimulus. When plasma glucose levels increase after food ingestion, uptake of glucose via glucose transporters (GLUT1 in humans and GLUT2 in rodents) [4,5] increases simultaneously, eventually resulting in the secretion of insulin. The canonical secretion pathway model describes that immediately after glucose uptake, glycolytic degradation of glucose to pyruvate results in enhanced oxidative phosphorylation, generating ATP and reducing the level of ADP in the cell. This elevation in the intracellular ATP/ADP ratio causes ATP-dependent potassium (K_ATP_) channels to close and subsequent depolarization of the membrane [6]. In turn, voltage-dependent Ca²⁺ channels are activated, the intracellular Ca²⁺ concentration rises, and exocytosis of insulin-filled vesicles is triggered, releasing the hormone into the bloodstream [7]. The following amplifying phase is K_ATP_ channel-independent [8]. By augmenting the response to increased intracellular Ca²^+^ concentrations, it allows for a reduced yet steady insulin secretion throughout the post-absorptive phase of a meal. Although studies with a focus on electrochemical characterization have dominated pancreatic beta cell research during the last decades, this canonical model has been challenged recently [9–11], and the potential role of additional mechanisms has been summarized in several reviews [12–14].

To gain a comprehensive understanding of the various levels of events underlying GSIS, transcriptomic, metabolomic, and phosphoproteomic approaches have been used to study the initial response of pancreatic beta cells to glucose stimulation [15–19]. The strength of these studies lies in their ability to capture changes throughout the whole cell. Particularly the metabolome and the phosphoproteome react very fast to environmental changes and are therefore perfect targets to investigate the initial response of pancreatic beta cells to a glucose stimulus, even for remarkably short incubation times, some as brief as 2 min [16,19]. However, many cellular processes depend directly on the regulation of protein levels, making the proteome response a crucial factor in insulin secretion mechanisms. Previous proteomic studies on beta cells focused on the influence of long-term exposure to compounds that trigger insulin secretion, ranging from 24 to 72 h [20–24]. In contrast, short-term changes of the proteome upon glucose stimulation remain so far uninvestigated, partly because of technical challenges. In this study, we investigated the proteomes of beta cells after 30-minute exposure to varying levels of glucose, ranging from glucose starvation up to very high glucose concentrations corresponding to blood glucose levels present in diabetic patients.

## Methods

### Reagents

| **Cell line** | **Vendor** | **Cat.-No.** |
| --- | --- | --- |
| INS-1 832/13 | Merck | SCC207 |

| **Assay/Plates** | **Vendor** | **Cat.-No.** |
| --- | --- | --- |
| Rat Insulin ELISA | Mercodia | 10-1250-01 |
| BCA 660 nm assay | Pierce™ | 22600 |
| BioPureSPE Macro- 96-well<br>Proto 300 C18 | The Nest Group | HNS S18V-L |
| Quantitative fluorometric<br>peptide assay | Pierce™ | 23290 |

| **Reagent** | **Vendor** | **Cat.-No.** |
| --- | --- | --- |
| RPMI 1640 | Sigma-Aldrich | R0885 |
| Fetal calf serum | Biochrom AG | S0115 |
| HEPES | Gibco | 15630-056 |
| glutamine | Millipore | K0283-BC |
| sodium-pyruvate | Millipore | TMS-005-C |
| penicillin | Millipore | A2213 |
| streptomycin | Millipore | A2213 |
| beta-mercaptoethanol | Thermo Fisher | 21985023 |
| glucose-free RPMI 1640 medium | Gibco | 11879020 |
| glucose | Gibco | A2494001 |
| urea | Sigma-Aldrich | U5128 |
| ammonium bicarbonate | Carl-Roth | T871.2 |
| magnesium chloride | AppliChem | 131396 |
| benzonase | Millipore | 101654 |
| dithiothreitol | Sigma-Aldrich | 43816 |
| iodoacetamide | Sigma-Aldrich | I1149 |
| trypsin Lys-C | Promega | V5072 |
| Formic acid | Fisher Scientific | A117-50 |
| methanol | Fisher Scientific | 10031094 |
| acetonitrile | Fisher Scientific | 10489553 |

### Cell culture

INS-1 832/13 cells were cultured in RPMI 1640 medium containing 11 mM glucose and supplemented with 10% fetal calf serum (FCS), 10 mM HEPES, 2 mM glutamine, 1 mM sodium-pyruvate, 50 mg penicillin, 50 mg streptomycin and 1x beta-mercaptoethanol. Cells were incubated at 37 °C in a humidified atmosphere at 5% CO_2_. Medium was exchanged every two days, three times a week. Cells were passaged weekly with a split ratio of 1:20.

### Glucose treatment and sample harvest

Cells were seeded in 12-well plates at 0.5 * 10⁶ cells/well. After 20 h of incubation, cells were washed twice with 1 mL of glucose-free RPMI 1640 medium supplemented with 10% FCS, 10 mM HEPES, 1 mM sodium pyruvate, 50 mg penicillin, 50 mg streptomycin and 1x beta-mercaptoethanol. Afterwards, the medium was replaced with 1 mL of RPMI 1640 supplemented with 3 mM glucose and cells were incubated 37 °C, 5% CO_2_ overnight to equilibrate to the lower glucose concentration. After overnight pre-incubation at 3 mM glucose, cells were washed twice with 1 mL glucose-free medium before 1 mL of stimulation medium (RPMI 1640 supplemented with 0, 2, 4, 5, 6, 7, 9, 10, 12, 15, or 20 mM glucose) was added. After adding the stimulation medium, cells were incubated for 30 min. The supernatant of each well was centrifuged at 400 x g, 4 °C for 5 min, and immediately assayed by ELISA in a 1:5 dilution following the manufacturer’s instructions. The cells were carefully washed twice on-plate with the respective stimulation medium before they were lysed on the plate using 100 μL of lysis buffer (8 M urea, 50 mM ammonium bicarbonate, 2 mM magnesium chloride, and freshly added 100 U benzonase) to extract proteins. Cell lysates were transferred to Eppendorf tubes and centrifuged at 16000 x g, 4 °C for 10 min. Supernatants were stored at −80 °C until further processing.

### Proteomics sample preparation

The protein concentration of lysates was quantified using a commercial bicinchoninic acid (BCA) 660 nm assay according to the manufacturer’s instructions. 50 μg of protein were processed per sample (in 8 M urea, 0.1 M ammonium bicarbonate). To reduce disulfide bonds, 5 mM dithiothreitol (final concentration) was added to each sample, followed by an incubation at 30 °C for 1 h. Next, 5 µL of 0.1 M iodoacetamide were added to each sample to acetylate previously reduced thiol groups. After an incubation of 30 min at 20 °C in the dark, samples were diluted by adding 340 µL of 0.1 M ABC before digestion with 1.5 µg of trypsin/LysC per sample. Samples were incubated overnight at 37 °C. Digestion was stopped by addition of 20 µL of 20% formic acid (FA). Peptides were purified by solid-phase extraction (SPE). SPE plates were activated by adding 200 μL/well of methanol. After centrifugation (50 x g for 1 min), 200 μL/well of 50% acetonitrile (ACN) were added, followed by another centrifugation step (150 x g for 1 min). The addition of ACN and the following centrifugation was repeated once. The flow-through was discarded and 200 μL/well of 0.1% FA were added. The plate was then centrifuged again (150 x g, 1 min). This step was repeated once before the flow through was discarded and 400 µL/well of the digested sample were added in two steps. After centrifugation (150 x g, 1 min), 200 μL/well of 0.1% FA was added, followed by another centrifugation (150 x g, 1 min). The addition of FA was repeated three times. A collection plate was placed underneath the SPE plate and the samples were eluted by three steps of addition of 110 μL of 50% ACN followed by centrifugation (200 x g, 1 min). The eluted samples were then dried in a vacuum concentrator and subsequently resuspended in 60 μL of 0.1% FA. To determine the peptide concentrations of the samples, a quantitative fluorometric peptide assay was carried out following the manufacturer’s protocol. Six process quality controls were prepared by pooling all samples, and then distributing this pooled sample once per row on the plate before peptide purification. Technical quality controls consisted of repeat injections of a single purified process quality control sample every ten measurements, resulting in 6 technical controls.

### LC-MS/MS measurements

Peptide separation was performed on an Ultimate 3000 RSLnanoHPLC (ThermoFisher Scientific). Tryptic peptides (1.25 μg/sample) were loaded on a trap column (PepMap C18, 5 mm x 300 μm x 5 μm, 100 Ǻ, Thermo Fisher Scientific) and separated on an analytical LC column (Acclaim PepMap C18, 200 mm x 75 μm x 2 μm; 100 Å; Thermo Fisher Scientific) at a flow rate of 250 nL/min. The mobile phase consisted of 0.1% FA (buffer A) and 80% ACN, 0.1% FA (buffer B). Digestion products were resolved using a linear gradient (3 - 20% buffer B in 80 min, followed by 20 - 45% buffer B in 10 min and 98% buffer B in 1 min). Total acquisition time per sample was 115 min. The eluate was directed to a Q Exactive Plus Hybrid Quadrupole-Orbitrap mass spectrometer (ThermoFisher Scientific), which operated in centroid mode. One MS1 scan was conducted at 35k resolving power with a maximum injection time of 60 ms, followed by 51 MS2 scans at 17.5k resolving power with a maximum injection time of 60 ms. The window length for the MS2 scans was set to a mass-to-charge ratio (m/z) of 24.0 using an overlapping window pattern. Samples were measured in data-independent acquisition mode and analyzed with the software DIA-NN [25]. Every tenth injection was a technical QC sample. Additionally, the six process QCs samples were randomly injected during the measurements.

### Pre-processing of proteomics data

Raw proteomic data were processed using DIA-NN 1.8 (https://github.com/vdemichev/DiaNN) [25] using the default settings with fragment ion m/z range set from 300-1800, mass accuracy set to 20 ppm at the MS2 and 10 ppm at the MS1 level, respectively, scan window set to 11, MBR (Match-between-runs) enabled, and the Quantification strategy set as “Robust LC (high Precision)”. To correctly assign bovine protein contaminations stemming from the use of FCS in the cultivation and wash medium, two FASTA files were provided for the generation of an in-silico spectral library and gene annotation: a Bos taurus reference proteome (UniProt UP000009136, accessed 2nd May 2022), and a Rattus Norvegicus reference proteome (UniProt UP000002494, accessed 15th September 2021). Species-specific gene interference was enabled. The report.tsv file resulting from the DIA-NN run was further processed at the precursor level using normalized precursor intensities. First, all non-proteotypic precursors and precursors with PG.Q.Value, Global.Q.Value, Global.PG.Q.Value, GG.Q.Value > 0.01, and precursors exhibiting a CV > 30% across the technical QCs (repeat injection) were removed. Only genes quantified by >=2 precursors, and genes quantified by exactly one precursor but detected in at least 4/6 replicates in every of the eleven conditions were retained, and protein quantification was performed using the diann_maxlfq() function of the DIA-NN R package (https://github.com/vdemichev/diann-rpackage). Bovine proteins were filtered out for all further analyses.

### Statistical analysis

#### Clustering analysis

For the cluster analysis, an ensemble clustering approach was adapted as described in [26]. For the ensemble clustering, the spatial *kMeans(++)*-[27,28], the density-based *commonNN*-[29–32], as well as the community-detection *Leiden*-algorithm [33] were combined. Since the *kMeans*-as well as the *commonNN*-algorithms are parameter-dependent, different parameter sets were included. For the *kMeans*-algorithms, the number of clusters *k* was varied between 10 and 49 in steps of 1 (40 different cluster results). For the *commonNN*-algorithm, the number of nearest neighbors in common *N* was varied between 2 and 10 in steps of 1, the distance cutoff R between 0.7\**R_cut* and 1.1\**R_cut* in steps of 0.1\**R_cut*, where *R_cut* denotes the distance at which roughly 0.1 % of all distances between the samples are smaller (45 different parameter combinations). The minimal number of samples per cluster *M* was fixed to 5. For the *Leiden* clustering, the seed was fixed to 42 and a directed 1-scaled(distance)-weighted network with *n_neighbors* = 10 was used, utilizing the *ModularityVertexPartition* algorithm.

For each algorithm, a separate co-clustering matrix was constructed and normalized by the number of different cluster results, where each element accounts for the probability that two samples were clustered together. The three co-clustering matrices were combined with equal weights and clustered again using a hierarchical Ward-clustering [34]. The clusters were extracted using a dynamic linkage-based cutoff. Due to the used algorithms, a block-diagonal matrix was obtained which was mainly driven by the Leiden-clustering, whereas the other two algorithms provided the fine-structure. In the end, 11 blocks with in total 104 subclusters were extracted. The analysis was carried out using *Python 3.9.13*, *numpy 1.22.4* [35], *scikit-learn 1.1.1* [36], *igraph 0.9.9* [37], *leidenalg 0.8.9* and *scipy 1.8.1* [38]. The used functions are available under https://github.com/OliverLemke/ensemble_clustering.

#### Over-representation analysis (ORA)

ORA was carried out using WebGestalt 2019 [39] using default settings. A list of all 3,703 consistently quantified proteins was used as the reference gene list and results were filtered at a false discovery rate (FDR) < 0.05. Multiple hypothesis correction to adjust P-values was applied (Benjamini-Hochberg correction) [40].

#### Differential expression analysis

Differential expression analysis was carried out using the *limma* package in R [41]. Linear models were fitted by the *lmFit()* function. Subsequently, empirical Bayes moderation was applied using the *eBayes()* function, resulting in log_2_FC and associated *P*-values. Multiple hypothesis correction to adjust *P*-values was applied (Benjamini-Hochberg correction) [40].

## Results

### The INS1 beta cell proteome after short-term glucose exposure

To characterize the dynamics of the beta cell proteome during the triggering phase of GSIS, we recorded glucose concentration-dependent insulin secretion and changes in protein abundances after glucose stimulation of 30 min in the pancreatic beta cell line INS-1 832/13 (Fig. 1a). Cells were stimulated with eleven different glucose concentrations between 0 - 20 mM, covering glucose starvation (0 mM), the physiological resting blood glucose level (3 - 5 mM), postprandial blood glucose levels (7 - 8 mM), as well as very high glucose levels often used in long-term glucose exposure studies (20 mM). Prior to stimulation, cells were incubated overnight at 3 mM glucose to adapt to resting glucose levels.

**Figure 1.**
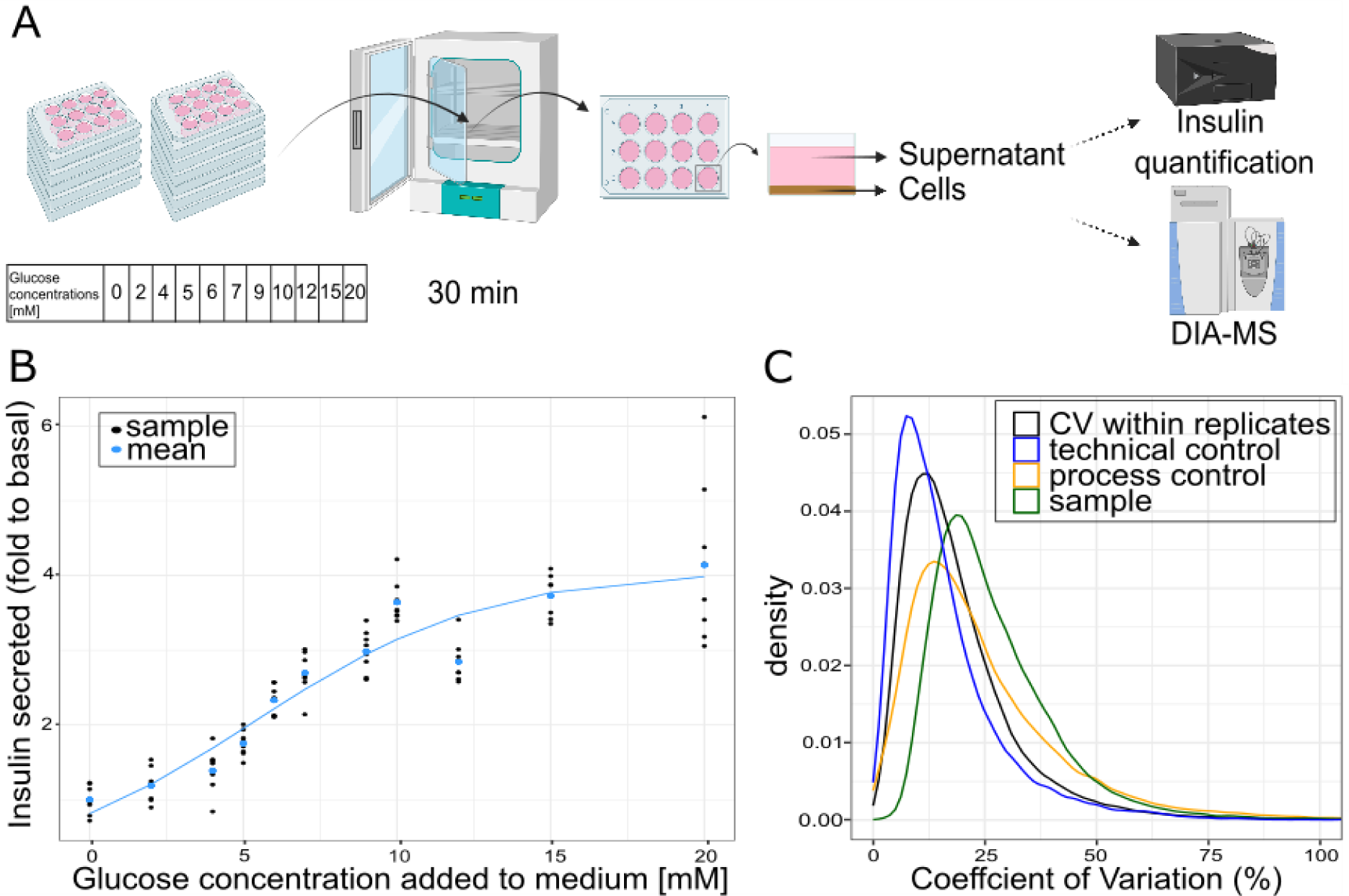
(A) Experimental design of the glucose stimulation dose response assay. INS-1 832/13 cells were cultivated in 12-well plates and each plate was treated with different glucose concentrations. In total, 11 concentrations in the range between 0 and 20 mM were used to stimulate cells for 30 min at 37°C, before the supernatant of all samples was used for insulin quantification by ELISA, and the cells were prepared for proteomic analysis. (B) Glucose concentration-dependent insulin secretion of INS-1 832/13 cells shown as fold change to basal (0 mM glucose) secretion. Individual replicates (n = 8) are shown in black dots and the mean of all replicates per condition is shown in blue dots. The data is fitted by the logistic function (blue line), with an residual standard error of 0.48 on 85 degrees of freedom. (C) Comparison of coefficients of variation (CV) within experimental groups (black) and across all biological samples (green). CVs within repeated technical control injections and sample preparation process controls are shown in blue and orange, respectively.

Insulin secretion by the cells was glucose concentration-dependent (Fig. 1b) and increased with glucose concentrations up to 10 mM glucose, where the amount of secreted insulin was around three times higher than without glucose stimulation (0 mM). The slope of the insulin secretion curve was flatter at lower glucose concentrations (0 to 4 mM), indicating less active, but not inactive, mechanisms of intracellular insulin secretion below this threshold. At glucose concentrations higher than 10 mM, insulin secretion stagnated.

Measuring glucose concentration-dependent changes in protein abundances upon short-term exposure poses technical challenges: most proteins are expected to show only small abundance changes, impeding the distinction of true biological responses from measurement noise. To mitigate these challenges, we measured the proteomes of six independently cultured replicates per condition and used a data-independent acquisition approach to increase the completeness of protein identifications across replicates and conditions. Before cell lysis, all samples were washed twice with the corresponding stimulation media to remove the extracellular proteome, including the secreted insulin. Wash and lysis steps were conducted on-plate to minimize sample handling time. The lysates were digested using trypsin and Lys-C overnight at 37°C, and resulting peptides were purified by solid-phase extraction. To be able to discriminate between biological changes in protein abundance in different stimulation conditions and technical variation inherent to the peptide purification process, we introduced six process controls (Materials and Methods). Furthermore, to monitor mass spectrometer performance over the run time and to estimate the technical variability of the measurements, we prepared another quality control pool that was injected every ten samples. The proteomic data was acquired using data-independent acquisition (DIA) with a 115 min gradient for each sample. Raw data were processed using DIA-NN [25] using an in-silico generated spectral library containing rat and bovine peptides to account for both proteins of interest expressed by the INS1 cells, as well as for potential contaminants present in the medium due to fetal calf serum addition. Only reliably detected precursors were retained for the quantification (Materials and Methods), and bovine proteins were excluded in subsequent analyses. In total, 3,703 rat proteins were consistently quantified across 66 samples (Table S1), with a very low fraction of missing values (0.31%). Measurement accuracy was very high, with the mode of the coefficient of variation (CV) among injection quality control samples at 7.78 % (Fig. 1c). The distributions of CVs within replicates of stimulation conditions and of the process quality control samples were comparable, with their modes at 11.75 % and 13.62 %, respectively. In contrast, the distribution of CVs calculated across all 66 samples was shifted towards higher values (mode at 19.35 %) compared to the control samples, indicating that despite the short glucose exposure time, we were able to record biological differences in the beta cell proteomes.

### Proteins show different response patterns to glucose stimulation

Next, we identified proteins with similar response profiles across different glucose stimulation conditions. For this, we applied an ensemble clustering approach as proposed and applied previously [26,42] by combining multiple distance-based cluster algorithms to account for differences at each time point. The resulting clusters (blocks) and subclusters of proteins denote similar concentration-dependent relative proteomic expression profiles (Fig. 2a, b). For the cluster analysis, we dropped the 0 mM glucose concentration data as it does not represent glucose stimulation, but, in contrast, glucose starvation. The 3,703 consistently measured proteins were grouped in 104 clusters in total, summarized in 11 blocks of clusters. Nine of these blocks contained proteins that showed only a weak response after short-term glucose stimulation, with median log_2_ fold changes (log_2_FC) not exceeding 0.4 and −0.4 at any condition. These blocks were further summarized into a single block encompassing 96 clusters containing in total 3,389 proteins (block 1, Fig. 2a). The second block included proteins that showed a slight increase of abundance upon glucose stimulation and a decrease at higher glucose concentrations (block 2, 5 clusters, 212 proteins, Fig. 2a, b). All protein clusters in that block show a specific curve shape, peaking at condition 6 mM with a median log_2_FC of 0.1 to 0.4, and reaching a stagnation level with a median log_2_FC of −0.5 at condition 15 mM. The third block contained proteins that followed the general pattern of block 2, but with much higher fold changes, peaking at 6 mM with median log_2_FC between 0.6 and 1.0. As in block 2, protein abundances decreased at higher glucose concentrations until returning to baseline expression levels between 0.2 and −0.1 at condition 15 mM (block 3, 3 clusters, 102 proteins, Fig. 2a, b).

**Figure 2.**
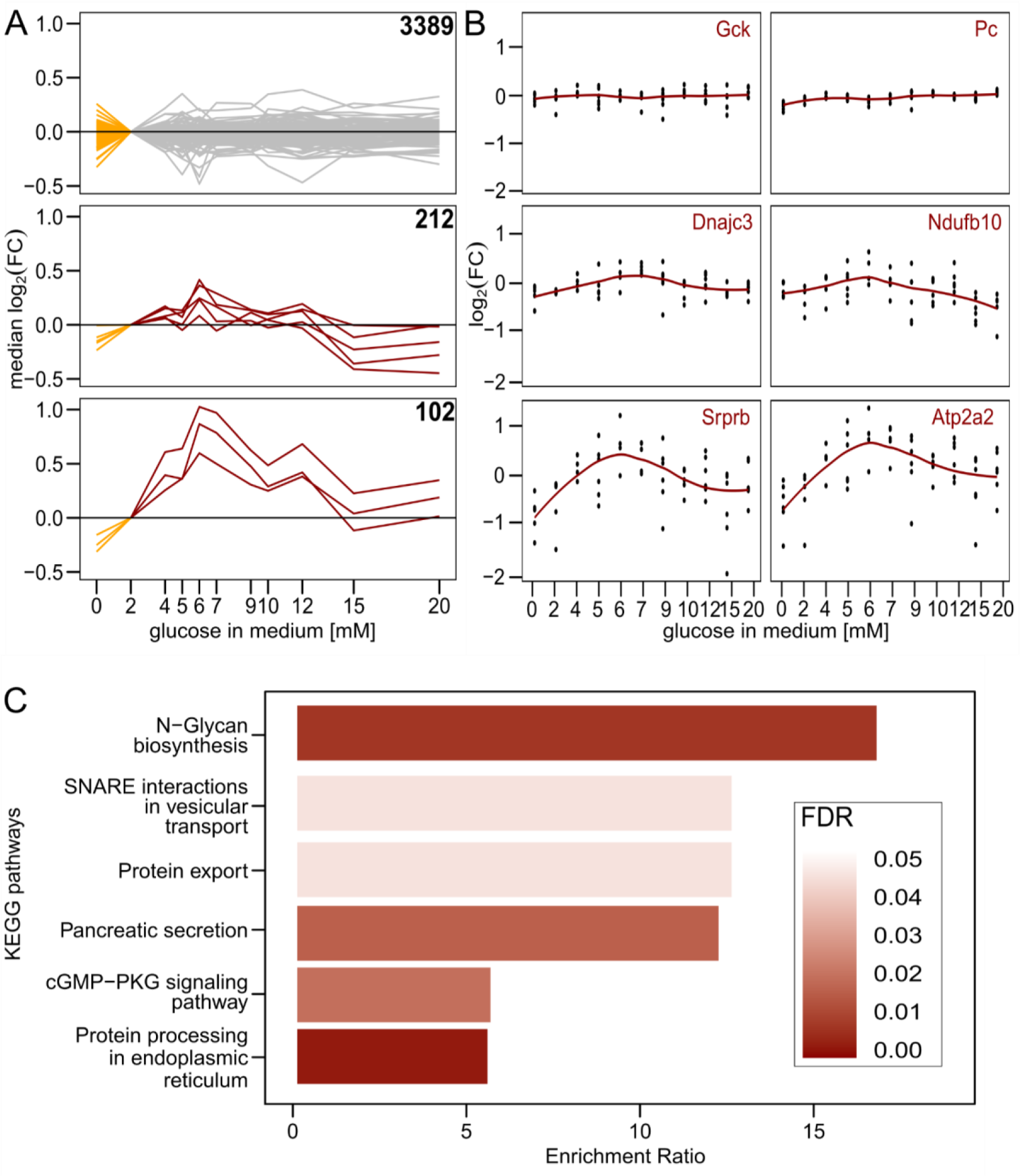
(A) Ensemble cluster analysis of the proteomic dataset. In total, 104 clusters were identified and similar clusters were further summarized into 3 different blocks (facets). Each line corresponds to one of the 104 clusters. The median of the log_2_ fold changes (log_2_FC) of the proteins in each cluster is shown. FCs were calculated relative to protein abundances at condition 2 mM. Gray protein clusters show little to no changes in protein abundance over the changing glucose concentration. Clusters that summarize proteins with changing abundances over the conditions are highlighted in red. Cluster analysis was performed for glucose stimulation conditions between 2 mM and 20 mM. The glucose starvation condition of 0 mM is depicted for each cluster in orange. (B) Abundance patterns of selected proteins that were assigned to block 1 (glucokinase (Gck), pyruvate carboxylase (Pc)), block 2 (DnaJ homolog subfamily C member three (Dnajc3), mitochondrial intermembrane space import and assembly protein 40 (Ndufb10)) or block 3 (signal recognition particle receptor subunit beta (Srprb), endoplasmic reticulum calcium ATPase 2 (Atp2a2)). Except for Gck and Pc in block 1, proteins are part of pathways that were enriched in the respective blocks. Each protein was measured in 6 replicates (black) of which the mean was calculated (red line). (C) Overrepresentation analysis for KEGG pathway terms for the 102 proteins of block 3. The false discovery rate (FDR) of each enriched pathway is indicated. Only significantly enriched pathways (FDR < 0.05) and pathways associated with pancreatic beta cells are shown (see also Table S4).

To determine whether certain gene sets were enriched in different protein blocks, we performed an overrepresentation analysis. All blocks that were summarized in block one were individually run in order to find specifically regulated pathways. However, because none of these protein blocks showed enriched pathways and due to the marginal abundance changes of proteins in block 1, we considered these proteins as not glucose concentration-dependently regulated. Interestingly, this block contained many proteins of major metabolic pathways. Important proteins of the glycolysis and the tricarboxylic acid (TCA) cycle which have been attributed a role in GSIS were found in this block, for example the glucokinase (GCK) and the pyruvate kinase (PK). Gck, the enzyme that phosphorylates glucose after entering the cell through the GLUT, showed no strong abundance change upon glucose stimulation with a maximum log_2_FC of 0.13 at 10 mM glucose. Parts of the protein complex of PK (Pklr and Pkm) were also identified in our data set, but not regulated in a glucose-concentration dependent manner (Pklr = maximum log_2_FC of 0.02 at 10 mM glucose, Pkm = maximum log_2_FC of 0.01 at 10 mM glucose). Furthermore, pyruvate carboxylase (PC), an enzyme that is critically linked to the TCA cycle as it helps replenish oxaloacetate, was also measured as not glucose-dependent regulated (Pc = maximum log_2_FC of 0.15 at 20 mM glucose). More proteins of these pathways were assigned to block 1, amongst them glucose-6-phosphate isomerase, phosphofructokinase, aldolase, triosephosphate isomerase 1, phosphoglycerate kinase 1, phosphoglycerate mutase 1, enolase, citrate synthase, isocitrate dehydrogenase, oxoglutarate dehydrogenase, succinate dehydrogenase, fumarate hydratase, malate dehydrogenase (Table S1). Interestingly, the main glucose uptake transporter in rodent beta cells (GLUT2) was, in contrast to the downstream glycolytic enzymes, sharply upregulated with increasing glucose concentrations and therefore appeared in block 3.

Block 2 revealed enrichment in several KEGG pathways, notably two associated with GSIS in pancreatic beta cells: protein processing in the endoplasmic reticulum (enrichment ratio = 4.5, FDR < 0.01) and oxidative phosphorylation (enrichment ratio = 3.1, FDR = 0.05) (Table S3). Similarly, Block 3 showed enrichment in multiple KEGG pathways (Fig. 2c, Table S4). Among the pathways with the highest enrichment ratio were several pathways that were expected to play a role in pancreatic beta cells after glucose stimulation, including SNARE interactions in vesicular transport and protein export (enrichment ratio = 10.8, FDR = 0.04), as well as pancreatic secretion (enrichment ratio = 10.5, FDR = 0.02) (Fig. 2c).

Next to glucose concentration-dependent upregulation of certain KEGG pathways, the analysis also revealed interesting protein responses to glucose starvation. While proteins from block 2 and 3 continuously show lower abundances (relative to 2 mM) at the starvation condition of 0 mM, certain protein clusters that are summarized in block 1 show higher protein abundances, unaltered abundances, or lower abundances at condition 0 (Fig. 2a, orange). Therefore, we performed a differential expression analysis between the conditions 0 mM and 2 mM, which identified 85 proteins as significantly upregulated (log_2_FC > 0.3, adj. p-Value < 0.05) and 15 proteins as significantly downregulated (log_2_FC < −0.3, p-Value < 0.05) among all proteins of the data set (Fig. 3a, Table S5). In the next step, overrepresentation analysis was performed on the up- and downregulated protein sets. While there was no enrichment within the downregulated proteins, for the upregulated proteins, the KEGG pathway ‘Fatty acid biosynthesis’ was identified (enrichment ratio = 23.4, FDR = 0.046). At the individual level, the abundance patterns of proteins assigned to this pathway, which were fatty acid synthase (FASN), acetyl CoA carboxylase 1 (ACCa, encoded by the Acaca gene), and long-chain fatty acid CoA ligase 1 and 4 (ACSL1 and ACSL4), showed a strong increase up to 4 mM of glucose (log_2_FC between 0.3 to 0.5 compared to condition 0 mM), followed by stagnation or slight decrease upon higher glucose concentrations (Fig. 3b).

**Figure 3.**
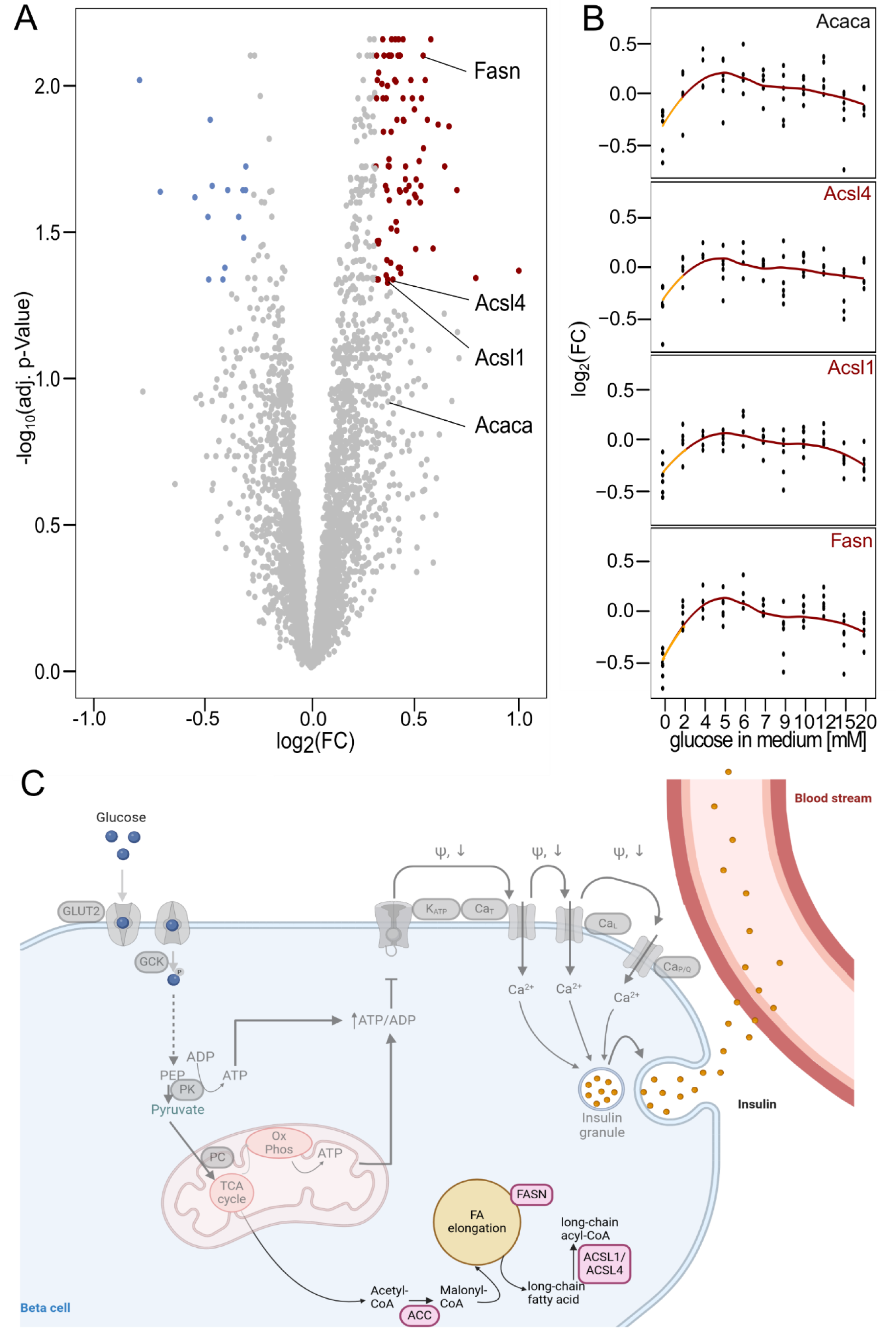
(A) Volcano plot showing differentially expressed proteins at 2 mM added glucose compared to the 0 mM glucose starvation condition. All significantly downregulated proteins (log_2_FC < −0.3, adj. p-Value < 0.05) are shown in blue, while all significantly upregulated proteins (log_2_FC > 0.3, adj. p-Value < 0.05) are shown in red. *P*-values were adjusted for multiple hypothesis testing using the Benjamini-Hochberg correction. The four annotated proteins belong to the fatty acid biosynthesis pathway, which was identified as the only enriched KEGG pathway among all significantly upregulated proteins. (B) Individual abundance patterns of the four proteins from (A). Each protein was measured in 6 replicates (black) of which the mean was calculated (red line). The orange line highlights the abundance change from 0 mM to 2 mM. (C) Illustration of proteins (light pink) that were identified as part of the overrepresented KEGG pathway fatty acid biosynthesis. The established GSIS model is indicated in transparent gray (Abbreviations: GLUT2 - glucose transporter 2, GCK - glucokinase, PEP - phosphoenolpyruvate, PK - pyruvate kinase, PC - pyruvate carboxylase, TCA cycle - tricarboxylic acid cycle, OxPhos - oxidative phosphorylation, ATP - adenosine triphosphate, ADP - adenosine diphosphate, ACC - acetyl CoA carboxylase, FASN - fatty acid synthase, ACSL - long chain fatty acid CoA ligase, FA - fatty acid, K_ATP_ - ATP dependent potassium channel, CA_T_ - t - type calcium channel, CA_L_ - l - type calcium channel, CA_P/Q_ - P/Q - type calcium channel, ω - membrane potential.

## Discussion

Glucose-stimulated insulin secretion has previously been intensively investigated at the metabolic [17,43,44] and electrochemical level [45–47]. Here, we characterized the behavior of the beta cell proteome during the triggering phase of GSIS by either stimulating INS-1 832/13 cells with varying glucose concentrations or starving the cells of glucose for 30 min. Using DIA proteomics, we obtained the dosage-dependent profiles of 3,703 proteins. We observed high variability across biological replicates, but the strength of our data set was notably bolstered by the inclusion of six biological replicates per condition, providing robust statistical power. The comprehensive replication gave us confidence in our ability to accurately measure even subtle changes in the proteome.

We were able to identify three different patterns of protein regulation by glucose concentration. The biggest group includes all proteins whose abundance is largely independent of glucose stimulation and starvation. The second pattern was caused by glucose concentration-dependent protein expression. The abundances of proteins in this group increased gradually up to a glucose concentration of 6 mM, and then decreased gradually at higher glucose concentrations. The third response pattern that we observed resembled a “switch-on” effect, characterized by a strong increase of abundance from 0 to 2 mM, followed by a slight increase at 4 mM and stagnation or a slight decrease of abundance at higher glucose concentrations. The identified proteins of group one included proteins that are prominently involved in the triggering phase of GSIS, such as individual enzymes of the glycolysis and the TCA cycle. This observation is consistent with a transcriptomic study that found that expression of genes associated with these processes is unaffected by rising extracellular glucose concentrations after 1 h of incubation with MIN6 cells [15]. The absence of upregulation of these metabolic pathways upon rising extracellular glucose concentration led the authors of the study to the conclusion that GSIS is independent of glucose consumption as a nutrient and that glucose instead might be more important as a signaling molecule [15].

A recent discussion about the possible necessity of a new model for glucose regulation in beta cells highlights the questions that still remain unanswered by the current model. The key point of debate was the relative importance of ATP generated by glycolysis versus oxidative phosphorylation [9–11]. Next to glycolytic enzymes, we were also able to identify enzymes from oxidative phosphorylation. The pathways showed up to be enriched among proteins that were assigned to block 2 during our cluster analysis, hence proteins that were slightly upregulated upon glucose concentrations until 6 mM. We therefore see that oxidative phosphorylation enzymes are slightly upregulated, while glycolysis enzymes are not. Other pathways that were expected to show up as being glucose concentration-dependently regulated were indeed enriched among the second protein group, for example pancreatic secretion, protein export, and SNARE interaction and vesicular transport.

Lastly, we identified proteins that are expressed upon release of glucose starvation: Differential expression analysis identified 85 proteins to be significantly upregulated at 2 mM compared to 0 mM glucose, among which only one KEGG pathway - Fatty acid biosynthesis - was enriched. A closer look into the abundance patterns of the proteins of this pathway revealed stagnation after a strong abundance increase at 2 - 4 mM, suggesting that this pathway might act in a switch-on pattern. The role of fatty acid biosynthesis in GSIS has been discussed in the literature [12,48,49], and malonyl-CoA, the product of ACC, has been suggested to be a regulative metabolic coupling factor [50].

A study by Lorenz et al. focusing on the short-term metabolic response of beta cells to glucose exposure [19] used a similar experimental design as herein, exposing INS 1 832/13 cells to 0-20 mM glucose for 30 min. The proteomic results presented here are in line with these metabolomic data, which revealed a strong increase in malonyl-CoA with a simultaneous decrease of acetyl-CoA. Interestingly, the abundance of malonyl-CoA is rising gradually with rising glucose concentrations [19]. In contrast, we observed the expression of ACCa to be switched on at low glucose concentrations but staying consistent over higher glucose concentrations. This supports the hypothesis that ACC acts as a coupling factor in beta cell metabolism through one of two mechanisms: either ACC is regulated at a level beyond protein expression, or with upregulated expression of the protein in the presence of low glucose concentrations, the maximum capacity of the enzyme is not reached even at high concentrations. The same trend extended to other proteins participating in fatty acid biosynthesis, such as FASN and ACSL, which exhibited similar abundance patterns as ACCa in our data with even stronger log_2_FC at 2 mM glucose.

Notably, previous studies have highlighted the significance of ACC, FASN, and ACSL4 in GSIS (Table 1). Inhibition of ACC and FASN by 5-(tetradecyloxy)-2-furoic acid (TOFA) and cerulenin, respectively, has demonstrated reduced GSIS induction in both rat pancreatic islets and INS-1 832/13 cells [51]. Additionally, several *in vitro* studies employing knock-out experiments in the INS-1 832/13 cell line have indicated that GSIS induction depends on the expression of ACC, FASN, or ACSL4 [52–55]. The results from the Acaca gene knock-out were further validated in an *in vivo* mice model [56].

**Table 1.**
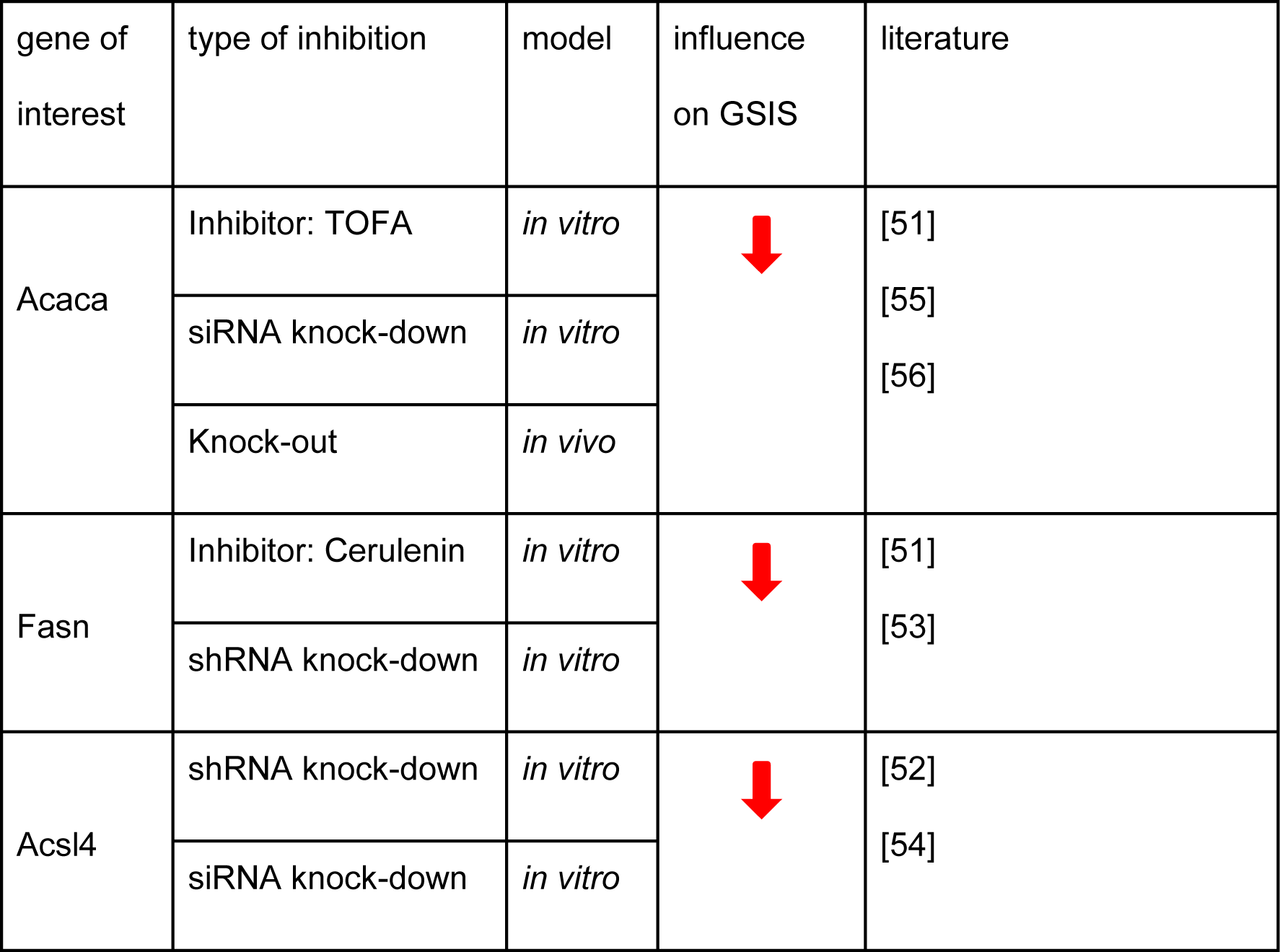
Impact of Fatty Acid Biosynthesis inhibition on GSIS.

Interestingly, the ongoing debate about glucose-regulated beta cell metabolism narrowly focuses on the generation of ATP to open K_ATP_ channels, while overlooking other biological processes that could potentially support GSIS [9–11]. Our data set, along with others, indicates the involvement of additional pathways immediately following glucose stimulation, such as fatty acid biosynthesis. But how could an increase in fatty acid metabolism support GSIS? We speculate about two possibilities: First, vesicle-mediated insulin secretion is dependent on membrane lipids that need to be replenished. Increased fatty acid biosynthesis might be required to support this process. Second, the increased glucose catabolism during GSIS creates a problem with electron flow. An increase in fatty acid biosynthesis might thus act as an electron acceptor. We hope our dataset stimulates further studies to investigate these possibilities.

### Associated Data and Code

Raw and processed mass spectrometry data have been deposited at the ProteomeXchange Consortium via the PRIDE partner repository with the dataset identifier PXD053750 and will be publicly released upon publication of the study. Functions used for clustering analysis are available at https://github.com/OliverLemke/ensemble_clustering.

### List of SI Tables

|  |  |
| --- | --- |
| Table S1 | Abundance ($\log_2$) of proteins of each replicate of all conditions and controls after preprocessing |
| Table S2 | Cluster and block assignment of individual proteins via ensemble clustering |
| Table S3 | Overrepresented KEGG pathways in block 2 of ensemble clustering |
| Table S4 | Overrepresented KEGG pathways in block 3 of ensemble clustering |
| Table S5 | Up- and downregulated proteins after differential expression analysis between 0 and 2 mM |

## Acknowledgements

We thank Sebastian Brachs, Anna Louise Gloyn, and Michele Solimena for useful discussions in the initial phase of the project, and Alexis Shih and Maike Sander for feedback in the final phase. Also, we thank the Core Facility High Throughput Mass Spectrometry of the Charité for support in sample preparation, acquisition, and analysis. Elements of figures were created with BioRender. This project has received funding from the Ministry of Education and Research (BMBF) as part of the National Research Node “Mass spectrometry in Systems Medicine” (MSCoreSys) under grant agreement 031L0220 (to M.R.) and from the European Research Council (ERC) under the European Union’s Horizon 2020 research and innovation programme (grant agreement No 951475 (to M.R.)).

M. R. is a founder and shareholder, and M. M. is a consultant and shareholder of Eliptica Ltd. All remaining authors declare no competing interests

## References

[1] IDF Diabetes Atlas 2021 n.d.

[2] Prentki, M., Matschinsky, F.M., Madiraju, S.R.M., Metabolic signaling in fuel-induced insulin secretion. Cell Metab. 2013, 18, 162–185.

[3] Huising, M.O., Paracrine regulation of insulin secretion. Diabetologia 2020, 63, 2057–2063.

[4] De Vos, A., Heimberg, H., Quartier, E., Huypens, P., et al., Human and rat beta cells differ in glucose transporter but not in glucokinase gene expression. J. Clin. Invest. 1995, 96, 2489–2495.

[5] McCulloch, L.J., van de Bunt, M., Braun, M., Frayn, K.N., et al., GLUT2 (SLC2A2) is not the principal glucose transporter in human pancreatic beta cells: implications for understanding genetic association signals at this locus. Mol. Genet. Metab. 2011, 104, 648–653.

[6] Nichols, C.G., KATP channels as molecular sensors of cellular metabolism. Nature 2006, 440, 470–476.

[7] Braun, M., The αβδ of ion channels in human islet cells. Islets 2009, 1, 160–162.

[8] Gembal, M., Detimary, P., Gilon, P., Gao, Z.Y., Henquin, J.C., Mechanisms by which glucose can control insulin release independently from its action on adenosine triphosphate-sensitive K+ channels in mouse B cells. J. Clin. Invest. 1993, 91, 871–880.

[9] Rutter, G.A., Sweet, I.R., Glucose Regulation of β-Cell KATP Channels: Is a New Model Needed? Diabetes 2024, 73, 849–855.

[10] Merrins, M.J., Kibbey, R.G., Glucose Regulation of β-Cell KATP Channels: It Is Time for a New Model! Diabetes 2024, 73, 856–863.

[11] Satin, L.S., Corradi, J., Sherman, A.S., Do We Need a New Hypothesis for KATP Closure in β-Cells? Distinguishing the Baby From the Bathwater. Diabetes 2024, 73, 844–848.

[12] Merrins, M.J., Corkey, B.E., Kibbey, R.G., Prentki, M., Metabolic cycles and signals for insulin secretion. Cell Metab. 2022, 34, 947–968.

[13] Campbell, J.E., Newgard, C.B., Mechanisms controlling pancreatic islet cell function in insulin secretion. Nat. Rev. Mol. Cell Biol. 2021, 22, 142–158.

[14] Kalwat, M.A., Cobb, M.H., Mechanisms of the amplifying pathway of insulin secretion in the β cell. Pharmacol. Ther. 2017, 179, 17–30.

[15] Firdos, Pramanik, T., Verma, P., Mittal, A., (Re-)Viewing Role of Intracellular Glucose Beyond Extracellular Regulation of Glucose-Stimulated Insulin Secretion by Pancreatic Cells. ACS Omega 2024, 9, 11755–11768.

[16] Sacco, F., Humphrey, S.J., Cox, J., Mischnik, M., et al., Glucose-regulated and drug-perturbed phosphoproteome reveals molecular mechanisms controlling insulin secretion. Nat. Commun. 2016, 7, 13250.

[17] Gooding, J.R., Jensen, M.V., Dai, X., Wenner, B.R., et al., Adenylosuccinate Is an Insulin Secretagogue Derived from Glucose-Induced Purine Metabolism. Cell Rep. 2015, 13, 157–167.

[18] Spégel, P., Sharoyko, V.V., Goehring, I., Danielsson, A.P.H., et al., Time-resolved metabolomics analysis of β-cells implicates the pentose phosphate pathway in the control of insulin release. Biochem. J 2013, 450, 595–605.

[19] Lorenz, M.A., El Azzouny, M.A., Kennedy, R.T., Burant, C.F., Metabolome Response to Glucose in the β-Cell Line INS-1 832/13. J. Biol. Chem. 2013, 288, 10923–10935.

[20] Waanders, L.F., Chwalek, K., Monetti, M., Kumar, C., et al., Quantitative proteomic analysis of single pancreatic islets. Proc. Natl. Acad. Sci. U. S. A. 2009, 106, 18902–18907.

[21] Pepaj, M., Bredahl, M.K., Gjerlaugsen, N., Bornstedt, M.E., Thorsby, P.M., Discovery of novel vitamin D-regulated proteins in INS-1 cells: a proteomic approach. Diabetes. Metab. Res. Rev. 2015, 31, 481–491.

[22] Kang, T., Jensen, P., Huang, H., Lund Christensen, G., et al., Characterization of the Molecular Mechanisms Underlying Glucose Stimulated Insulin Secretion from Isolated Pancreatic β-cells Using Post-translational Modification Specific Proteomics (PTMomics). Mol. Cell. Proteomics 2018, 17, 95–110.

[23] Li, Z., Liu, H., Niu, Z., Zhong, W., et al., Temporal Proteomic Analysis of Pancreatic β-Cells in Response to Lipotoxicity and Glucolipotoxicity*. Mol. Cell. Proteomics 2018, 17, 2119–2131.

[24] Lytrivi, M., Ghaddar, K., Lopes, M., Rosengren, V., et al., Combined transcriptome and proteome profiling of the pancreatic β-cell response to palmitate unveils key pathways of β-cell lipotoxicity. BMC Genomics 2020, 21, 590.

[25] Demichev, V., Messner, C.B., Vernardis, S.I., Lilley, K.S., Ralser, M., DIA-NN: neural networks and interference correction enable deep proteome coverage in high throughput. Nat. Methods 2020, 17, 41–44.

[26] Ronan, T., Qi, Z., Naegle, K.M., Avoiding common pitfalls when clustering biological data. Sci. Signal. 2016, 9, re6.

[27] Jin, X., Han, J., in:, Sammut C, Webb GI (Eds.), Encyclopedia of Machine Learning, Springer US, Boston, MA 2010, pp. 563-564.

[28] Arthur, D., Vassilvitskii, S., in:, Proceedings of the Eighteenth Annual ACM-SIAM Symposium on Discrete Algorithms, SODA 2007, New Orleans, Louisiana, USA, January 7-9, 2007, vol. 8, unknown, 2007, pp. 1027-1035.

[29] Keller, B., Daura, X., van Gunsteren, W.F., Comparing geometric and kinetic cluster algorithms for molecular simulation data. J. Chem. Phys. 2010, 132, 074110.

[30] Lemke, O., Keller, B.G., Density-based cluster algorithms for the identification of core sets. J. Chem. Phys. 2016, 145, 164104.

[31] Lemke, O., Keller, B.G., Common Nearest Neighbor Clustering—A Benchmark. Algorithms 2018, 11, 19.

[32] Kapp-Joswig, J.-O., Keller, B.G., CommonNNClustering─A Python Package for Generic Common-Nearest-Neighbor Clustering. J. Chem. Inf. Model. 2023, 63, 1093–1098.

[33] Traag, V.A., Waltman, L., van Eck, N.J., From Louvain to Leiden: guaranteeing well-connected communities. Sci. Rep. 2019, 9, 5233.

[34] Ward, J.H., Hierarchical Grouping to Optimize an Objective Function. J. Am. Stat. Assoc. 1963, 58, 236–244.

[35] Harris, C.R., Millman, K.J., van der Walt, S.J., Gommers, R., et al., Array programming with NumPy. Nature 2020, 585, 357–362.

[36] Pedregosa, F., Varoquaux, G., Gramfort, A., Michel, V., et al., Scikit-learn: Machine Learning in Python. J. Mach. Learn. Res. 2011, 12, 2825–2830.

[37] Horvát, S., Nepusz, T., Traag, V., Zanini, F., Csárdi, G., igraph — the network analysis package, 2020.

[38] Virtanen, P., Gommers, R., Oliphant, T.E., Haberland, M., et al., SciPy 1.0: fundamental algorithms for scientific computing in Python. Nat. Methods 2020, 17, 261–272.

[39] Liao, Y., Wang, J., Jaehnig, E.J., Shi, Z., Zhang, B., WebGestalt 2019: gene set analysis toolkit with revamped UIs and APIs. Nucleic Acids Res. 2019, 47, W199–W205.

[40] Klipper-Aurbach, Y., Wasserman, M., Braunspiegel-Weintrob, N., Borstein, D., et al., Mathematical formulae for the prediction of the residual beta cell function during the first two years of disease in children and adolescents with insulin-dependent diabetes mellitus. Med. Hypotheses 1995, 45, 486–490.

[41] Ritchie, M.E., Phipson, B., Wu, D., Hu, Y., et al., limma powers differential expression analyses for RNA-sequencing and microarray studies. Nucleic Acids Res. 2015, 43, e47.

[42] Aulakh, S.K., Lemke, O., Szyrwiel, L., Kamrad, S., et al., The molecular landscape of cellular metal ion biology. bioRxiv 2024, 2024.02.29.582718.

[43] Barillaro, M., Schuurman, M., Wang, R., β1-Integrin-A Key Player in Controlling Pancreatic Beta-Cell Insulin Secretion via Interplay With SNARE Proteins. Endocrinology 2022, 164.

[44] Wang, Z., Archang, M., Gurlo, T., Wong, E., et al., Application of fluorescence lifetime imaging microscopy to monitor glucose metabolism in pancreatic islets in vivo. Biomed. Opt. Express 2023, 14, 4170–4178.

[45] Misler, S., Barnett, D.W., Gillis, K.D., Pressel, D.M., Electrophysiology of stimulus-secretion coupling in human beta-cells. Diabetes 1992, 41, 1221–1228.

[46] Jacobson, D.A., Kuznetsov, A., Lopez, J.P., Kash, S., et al., Kv2.1 ablation alters glucose-induced islet electrical activity, enhancing insulin secretion. Cell Metab. 2007, 6, 229–235.

[47] Rorsman, P., Eliasson, L., Kanno, T., Zhang, Q., Gopel, S., Electrophysiology of pancreatic β-cells in intact mouse islets of Langerhans. Prog. Biophys. Mol. Biol. 2011, 107, 224–235.

[48] Corkey, B.E., Glennon, M.C., Chen, K.S., Deeney, J.T., et al., A role for malonyl-CoA in glucose-stimulated insulin secretion from clonal pancreatic beta-cells. J. Biol. Chem. 1989, 264, 21608–21612.

[49] Zhang, S., Kim, K.H., Essential role of acetyl-CoA carboxylase in the glucose-induced insulin secretion in a pancreatic beta-cell line. Cell. Signal. 1998, 10, 35–42.

[50] Prentki, M., Vischer, S., Glennon, M.C., Regazzi, R., et al., Malonyl-CoA and long chain acyl-CoA esters as metabolic coupling factors in nutrient-induced insulin secretion. J. Biol. Chem. 1992, 267, 5802–5810.

[51] MacDonald, M.J., Dobrzyn, A., Ntambi, J., Stoker, S.W., The role of rapid lipogenesis in insulin secretion: Insulin secretagogues acutely alter lipid composition of INS-1 832/13 cells. Arch. Biochem. Biophys. 2008, 470, 153–162.

[52] Ansari, I.-U.H., Longacre, M.J., Stoker, S.W., Kendrick, M.A., et al., Characterization of Acyl-CoA synthetase isoforms in pancreatic beta cells: Gene silencing shows participation of ACSL3 and ACSL4 in insulin secretion. Arch. Biochem. Biophys. 2017, 618, 32–43.

[53] MacDonald, M.J., Hasan, N.M., Dobrzyn, A., Stoker, S.W., et al., Knockdown of pyruvate carboxylase or fatty acid synthase lowers numerous lipids and glucose-stimulated insulin release in insulinoma cells. Arch. Biochem. Biophys. 2013, 532, 23–31.

[54] Klett, E.L., Chen, S., Edin, M.L., Li, L.O., et al., Diminished acyl-CoA synthetase isoform 4 activity in INS 832/13 cells reduces cellular epoxyeicosatrienoic acid levels and results in impaired glucose-stimulated insulin secretion. J. Biol. Chem. 2013, 288, 21618–21629.

[55] Ronnebaum, S.M., Joseph, J.W., Ilkayeva, O., Burgess, S.C., et al., Chronic suppression of acetyl-CoA carboxylase 1 in beta-cells impairs insulin secretion via inhibition of glucose rather than lipid metabolism. J. Biol. Chem. 2008, 283, 14248–14256.

[56] Cantley, J., Davenport, A., Vetterli, L., Nemes, N.J., et al., Disruption of beta cell acetyl-CoA carboxylase-1 in mice impairs insulin secretion and beta cell mass. Diabetologia 2019, 62, 99–111.

